# Comparison of SARS-CoV-2 infections among 3 species of non-human primates

**DOI:** 10.1101/2020.04.08.031807

**Authors:** Shuaiyao Lu, Yuan Zhao, Wenhai Yu, Yun Yang, Jiahong Gao, Junbin Wang, Dexuan Kuang, Mengli Yang, Jing Yang, Chunxia Ma, Jingwen Xu, Xingli Qian, Haiyan Li, Siwen Zhao, Jingmei Li, Haixuan Wang, Haiting Long, Jingxian Zhou, Fangyu Luo, Kaiyun Ding, Daoju Wu, Yong Zhang, Yinliang Dong, Yuqin Liu, Yingqiu Zheng, Xiaochen Lin, Li Jiao, Huanying Zheng, Qing Dai, Qiangmin Sun, Yunzhang Hu, Changwen Ke, Hongqi Liu, Xiaozhong Peng

**Affiliations:** National Kunming High-level Biosafety Primate Research Center, Institute of Medical Biology, Chinese Academy of Medical Sciences and Peking Union Medical College, Yunnan China; State Key Laboratory of Medical Molecular Biology, Department of Molecular Biology and Biochemistry, Institute of Basic Medical Sciences, Medical Primate Research Center, Neuroscience Center, Chinese Academy of Medical Sciences, School of Basic Medicine Peking Union Medical College, Beijing China; Medical Key Laboratory for Repository and Application of Pathogenic Microbiology, Guangdong Provincial Center for Disease Control and Prevention, Guangzhou China

## Abstract

COVID-19, caused by SARS-CoV-2 infection, has recently been announced as a pandemic all over the world. Plenty of diagnostic, preventive and therapeutic knowledges have been enriched from clinical studies since December 2019. However, animal models, particularly non-human primate models, are urgently needed for critical questions that could not be answered in clinical patients, evaluations of anti-viral drugs and vaccines. In this study, two families of non-human primates, Old world monkeys (12 *Macaca mulatta*, 6 *Macaca fascicularis*) and New world monkeys (6 *Callithrix jacchus*), were experimentally inoculated with SARS-CoV-2. Clinical signs were recorded. Samples were collected for analysis of viral shedding, viremia and histopathological examination. Increased body temperature was observed in 100% (12/12) *M. mulatta*, 33.3% (2/6) *M. fascicularis* and none (0/6) of *C. jacchus* post inoculation of SARS-CoV-2. All of *M. mulatta* and *M. fascicularis* showed chest radiographic abnormality. Viral genomes were detected in nasal swabs, throat swabs, anal swabs and blood from all 3 species of monkeys. Viral shedding from upper respiratory samples reached the peak between day 6 and day 8 post inoculation. From necropsied *M. mulatta* and *M. fascicularis*, the tissues showing virus positive were mainly lung, weasand, bronchus and spleen. No viral genome was seen in any of tissues from 2 necropsied *C. jacchus.* Severe gross lesions and histopathological changes were observed in lung, heart and stomach of SARS-CoV-2 infected animals. In summary, we have established a NHP model for COVID-19, which could be used to evaluate drugs and vaccines, and investigate viral pathogenesis. *M. mulatta* is the most susceptible to SARS-CoV-2 infection, followed by *M. fascicularis* and *C. jacchus.*

**One Sentence Summary:** *M. mulatta* is the most susceptible to SARS-CoV-2 infection as compared to *M. fascicularis* and *C. jacchus*.

## INTRODUCTION

The novel coronavirus disease COVID-19, caused by SARS-CoV-2 infection, was firstly reported in Wuhan, Hubei China in December 2019^1^. About one month, this severe disease attracted international attentions and declared as a Public Health Emergency of International Concern (PHEIC) on January 30, 2020. Within 4 months, COVID-19 unexpectedly spread to 195 countries and regions, led to more than 1 million confirmed cases in the world and was defined as a pandemic disease. It is believed to be the second largest infectious disease since 1918 flu^2^. Clinicians and scientists have been globally collaborating to investigate this disease and its pathogen, such as diagnosis, prevention and treatment of COVID-19^3^, genome and structure of SARS-CoV-2^4^. However, there are still many critical questions remaining, which are hard to be answered in clinic studies. For example, how do SARS-CoV-2 transmit? What is viral pathogenesis? What about vaccines and drugs against SARS-CoV-2? Therefore, animal models are urgently expected to be established for this severe COVID-19. Although several models of COVID-19 have recently been reported^5-7^, including the non-human primate model, none of model should be considered to be the best one, depending on what questions we are going to answer with the animal models. About 16 years ago, the non-human primate models recapitulated several important aspects of SARS ^8-11^. These models made great contributions to investigation of SARS pathogenesis, evaluation of antiviral drugs and vaccine ^12^. Though COVID-19 is different from SARS in some aspects, pathogens for these two diseases share some characters, such as ACE2 receptor. Here, to establish the COVID-19 model, two families including 3 species of non-human primates, which are widely used for animal models with their own advantages and disadvantages, were experimentally infected with SARS-CoV-2, followed by comparisons of clinical symptoms, hematology, biochemical indexes, immunology and histopathology among 3 species. We found that both *Macaca mulatta (M. mulatta)* and *Macaca fascicularis* (*M. fascicularis*) of old world monkeys were susceptible to SARS-CoV-2 infection, although *M. mulatta* was more susceptible to viral infection than *M. fascicularis*. However, *Callithrix jacchus* (*C. jacchus*), a new world monkey, was relatively resistant to SARS-CoV-2 infection. Therefore, as the model of COVID-19, *M. mulatta* and *M. fascicularis* may be chosen for further exploration of viral pathogenesis, evaluation of anti-SARS-CoV-2 drugs and vaccines.

## RESULTS

Given that host factors may be involved in viral pathogenesis, we designed an experiment in the present study to investigate whether host genetics, age and gender affect SARS-CoV-2 infection in non-human primates (Figure 1). Two families of non-human primates, old world monkeys (12 *M. mulatta*, 6 *M. fascicularis*) and new world monkeys (6 *C. jacchus*), were chosen for this experiment after screening and randomly grouped based on species, age and gender (Figure 1). Animals in large size (4 adults and 4 old *M. mulatta*, 6 *M. fascicularis*) were inoculated with 4.75×10^6^ pfu of SARS-CoV-2 via 3 routines (intratreachuslly 4.0ml, intranasally 0.5ml and intra 0.25ml). Half dosage of the viruses was given to 4 young *M. mulatta* via the same 3 routines. Six *C. jacchus* were inoculated with 1.0×10^6^ pfu intranasally.

**Figure 1.**
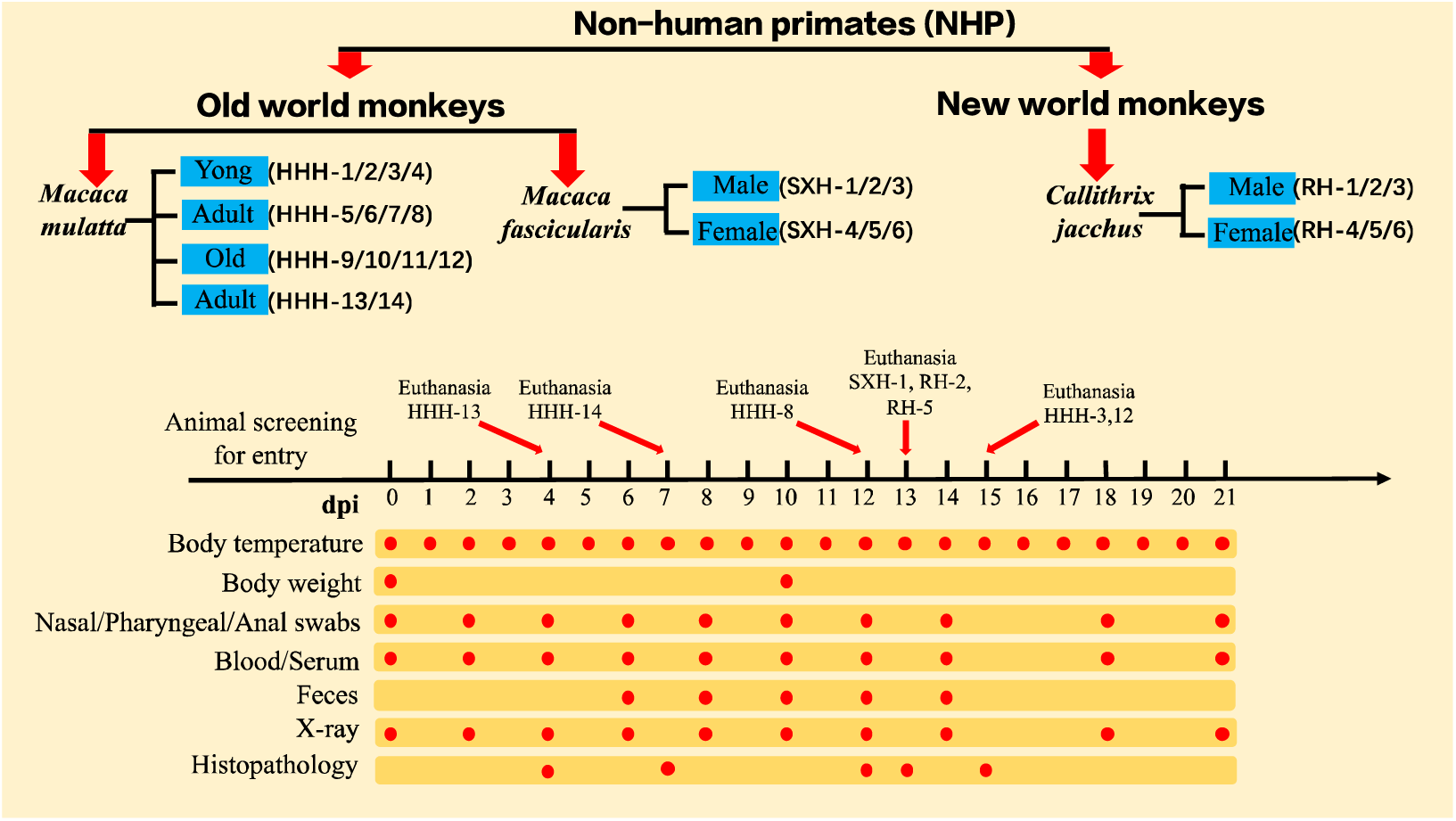
Schematic of the study design. Two families of non-human primates including 3 species of monkeys (totally 26 animals) were selected for this comparative study of modelling COVID-19. Age and gender were considered for grouping monkeys. After collection of baseline samples, all animals were inoculated with SARS-CoV-2 as stated in Materials and Methods. Clinical signs, virus shedding and replication, host responses to SARS-CoV-2 were recorded and evaluated at the indicated time points.

### Clinical signs of SARS-CoV-2 infected monkeys

Animals were monitored daily and sampled at the indicated time points before and after viral inoculation (Figure 1). Increased body temperature (BT) (above 38 °C) was continuously observed in 100% (12/12) *M. mulatta*, 33.3% (2/6) *M. fascicularis* and none (0/6) of *C. jacchus* post inoculation of SARS-CoV-2. BT of 7/12 *M. mulatta* reached the first peak on day 4-6 post inoculation (dpi) and maintained above 38 °C for several days. The highest BT was 40.9 °C during SARS-CoV-2 infection. Of *M. fascicularis*, their BT didn’t change as much as those of *M. mulatta*. And duration of BT>38 °C in *M. fascicularis* was shorter than *M. mulatta* (Figure 2A).

**Figure 2.**
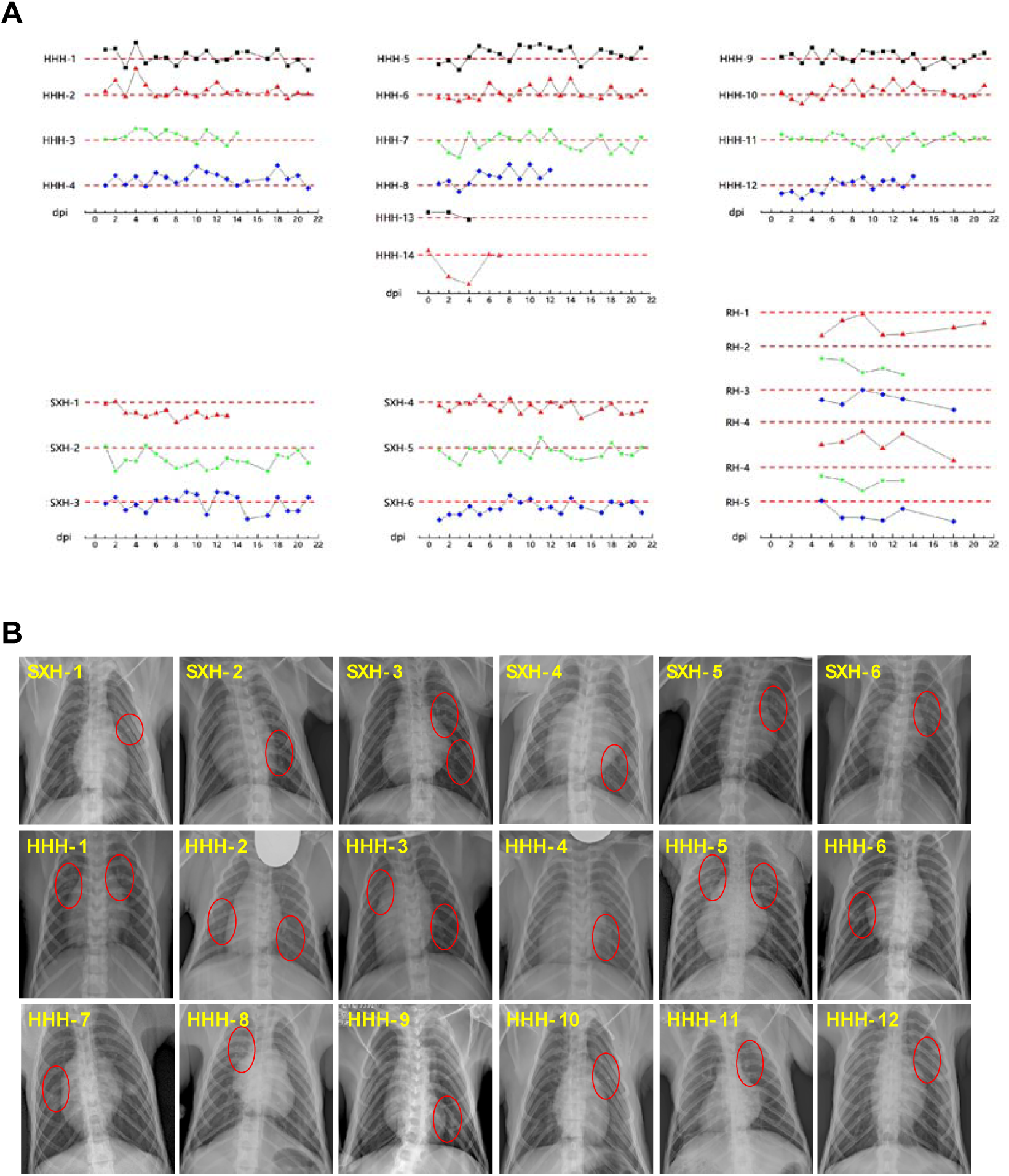
Clinical signs of SARS-CoV-2 infected monkeys. (A) Body temperature of monkeys was monitored and recorded every day after SARS-CoV-2 inoculation. Change curves of body temperature were prepared via OriginLab software. (B) Chest radiograph of SARS-CoV-2 challenged monkeys was recorded every two days by Mobile digital medical X-ray photography system. Radiographs were graded by two thoracic radiologists independently and double-blindly via a scoring system published for MERS and influenza as described in Methods.

On 0 and 10 dpi, body weight (BW) of experimental animals (*C. jacchus* not determined) was checked on electronic balance. As compared to BW on day 0, nine out of 12 *M. mulatta* showed the decreased BW with declined range of 5.88%-28.57%. Eighty-three percent (5/6) of *M. fascicularis* lost BW by 2.17-10.51% (Table 1).

**Table 1.** Body weight changes of SARS-CoV-2 infected monkeys

| Species | Groups | Monkey ID | Body weight changes* |
| --- | --- | --- | --- |
| *Macaca fascicularis* | Male | SXH-1 | 2.17% ↓ |
|  |  | SXH-2 | 2.34% ↓ |
|  |  | SXH-3 | 10.51% ↓ |
|  | Female | SXH-4 | 2.31% ↓ |
|  |  | SXH-5 | 0.00% |
|  |  | SXH-6 | 6.67% ↓ |
| *Macaca mulatta* | Young | HHH-1 | 0.00% ↓ |
|  |  | HHH-2 | 5.88% ↑ |
|  |  | HHH-3 | 28.57% ↓ |
|  |  | HHH-4 | 0.00% |
|  | Adult | HHH-5 | 11.43% ↓ |
|  |  | HHH-6 | 10.77% ↓ |
|  |  | HHH-7 | 10.77% ↓ |
|  |  | HHH-8 | 15.12% ↓ |
|  | Old | HHH-9 | 13.04% ↓ |
|  |  | HHH-10 | 12.22% ↓ |
|  |  | HHH-11 | 7.72% ↓ |
|  |  | HHH-12 | 11.36% ↓ |

\*Body weights (BW) of animals were measured on day 2 and day 6 post virus inoculation. Body weight change (%) was expressed as $|\text{BW on 6 dpi} - \text{BW on 2 dpi}| / \text{BW on 2 dpi}$ .

Chest radiographs of infected animals (*M. mulatta* and *M. fascicularis*, not *C. jacchus* due to their small size) were recorded every other day post virus inoculation. From the beginning of 10 dpi, abnormalities at various degrees appeared in lungs of all *M. mulatta* and *M. fascicularis*. The texture of the two lungs is increased and thickened, and scattered in small patches (Figure 2B). Overview of the chest radiograph throughout this study showed that progressive pulmonary infiltration was noted in all *M. mulatta* and *M. fascicularis* (Supplemental Figure 1).

### Replication of SARS-CoV-2 in the non-human primates

To know dynamics of viral replication and virus shedding, samples of nasal swabs, throat swabs, anal swabs, feces, blood and tissues were collected at the indicated time points, and SARS-CoV-2 genomes were quantitated by RT-qPCR. Swab samples collected on 2 dpi from *M. mulatta* and *M. fascicularis* showed surprisingly high levels of viral genome RNA, particularly in nasal swabs. Most of swab samples saw the second peaks of viral RNA on 6-8 dpi. In some swab samples from old world monkeys, viral RNA was still detectable on 14 dpi (Figure 3A). Less viral RNA were detected in throat swabs, compared to nasal swabs and anal swabs. In contrast to *M. mulatta* and *M. fascicularis*, lower levels of viral RNA were detected in swab samples from *C. jacchus* during two weeks post viral inoculation (Figure 3B). Virus shedding in feces from 8 out of 18 old world monkeys started on 6 dpi. Notably, higher levels of viral RNA were detected in feces from SXH6, HHH6, HHH11 and HHH12, from which anal swabs also gave correspondingly higher number of viral RNA. In peripheral blood, 8/18 old world monkeys and 6/6 new world monkeys had viral RNA detectable on 2 dpi. After 6 days, blood samples from nearly all old world monkeys became viral RNA-positive. On 10 dpi, viral RNA was not detectable in blood samples from almost all (23/24) of experimental monkeys (Figure 3A).

**Figure 3.**
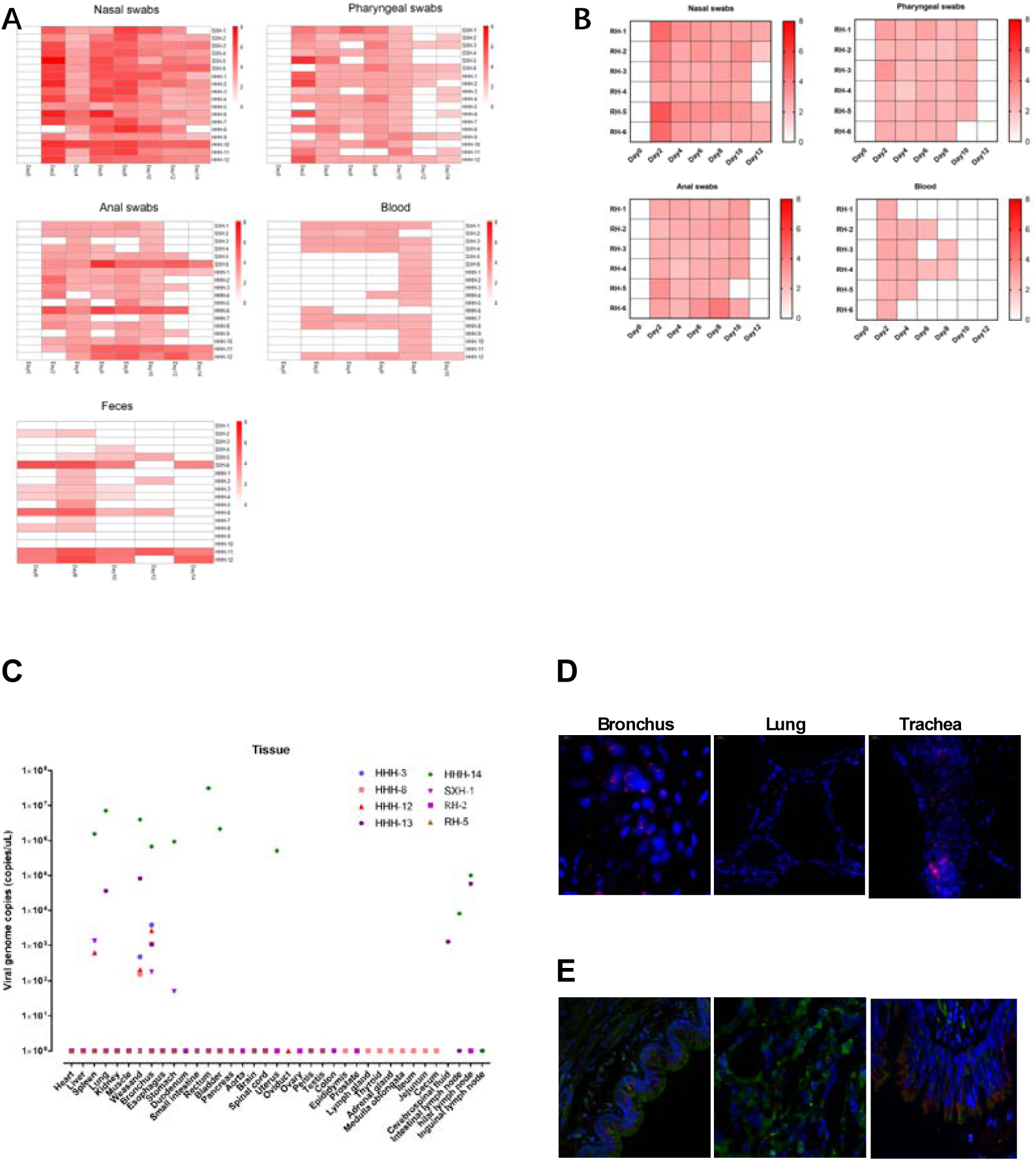
Virus shedding and replication of SARS-CoV-2 in the NHP. Every other day post virus inoculation, swab samples (from nasal, throat, anal) and blood samples of monkeys were collected for quantification of virus genomic RNA via RT-qPCR (A. Old world monkeys; B. New world monkeys). Tissue samples were harvested from eight necropsied animals at the indicated time points for evaluation of viral loads by RT-qPCR (C). For viral localization in tissues, nucleic acid hybridization of RNAsope technology was performed for detection of viral RNA via specific probe for SARS-CoV-2 (D) and immunofluorescent assay was conducted to detect viral S antigen (red) and ACE2 (green) (E).

Viral load of SARS-CoV-2 was determined in tissue samples from 8 animals necropsied on 4 dpi (HHH13), 7 dpi (HHH14), 12 dpi (HHH8), 13 dpi (SXH-1, RH2, RH5) and 15 dpi (HHH3, HHH12). RT-qPCR with SARS-CoV-2 specific primers and probes was performed to quantitate copy number of viral genomic RNA. No viral RNA was detected from all tissues of two *C. jacchus*. We could detect higher level of viral genomes in the tissue of bronchus and weasand from 3 necropsied *M. mulatta* and relatively lower level of viral RNA in one *M. fascicularis*. Spleens from HHH12 and SXH1 gave middle levels of viral RNA. No viral RNA was detected in other tissues. Furthermore, viral RNA and spike protein in tissue samples were confirmed in lung, trachea and bronchus by RNA hybridization RNAscope and immunofluorescent assay (Figure 3D and Figure 3E). Under TEM, no viral particle was observed in the ultrathin sections of lungs and other tissues.

### Host responses to SARS-CoV-2 infection

To determine the host response to SARS-CoV-2 infection, we firstly performed comprehensive analyses of peripheral blood and serum samples collected at the indicated time points post infection. Blood complete test (BCT), blood clotting assay (BCA), flow cytometric assay (FCA) were conducted to evaluate the anti-coagulated whole blood harvested every other day post viral inoculation. Twenty-three parameters of whole blood in BCT were assayed, related to white blood cells (WBC), red blood cells (RBC) and platelets. BCT results showed a slight and irregular changes during infection, which indicated no clinical meaning. In BCA, three parameters of PT-R, PT-T, PT-INR progressively decreased with infection, suggesting that SARS-CoV-2 infection may affect blood coagulation function (data not shown). Three major populations of immune cells in RBC-depleted peripheral blood were differentially and dynamically analyzed via flow cytometry. In *M mulatta*, regardless of ages, frequencies of CD4+ T cells, CD8+ T cells, monocytes peaked on 2 or 4 dpi, and then gradually declined. Young *M mulatta* showed the strongest B cell responses to SARS-CoV-2 infection among 3 ages of monkeys, but they had the similarly changing trends (peaked on 2 or 4 dpi and maintained at a certain level) (Figure 4A). *M. fascicularis* showed the similar dynamics of CD4+ T cells, CD8+ T cells, B cells and monocytes in blood post SARS-CoV-2 infection, of which no significant difference between male and female animals (Figure 4B). In SARS-CoV-2 infected monkeys, serum samples were collected for assay of biochemistry indexes related with liver functions, heart/muscle functions, kidney functions, blood lipids, humoral immune responses, and stresses (data not shown). Unfortunately, we could not find any biochemistry index that could be a biomarker of SARS-CoV-2 infection correlated with functions of key organs.

**Figure 4.**
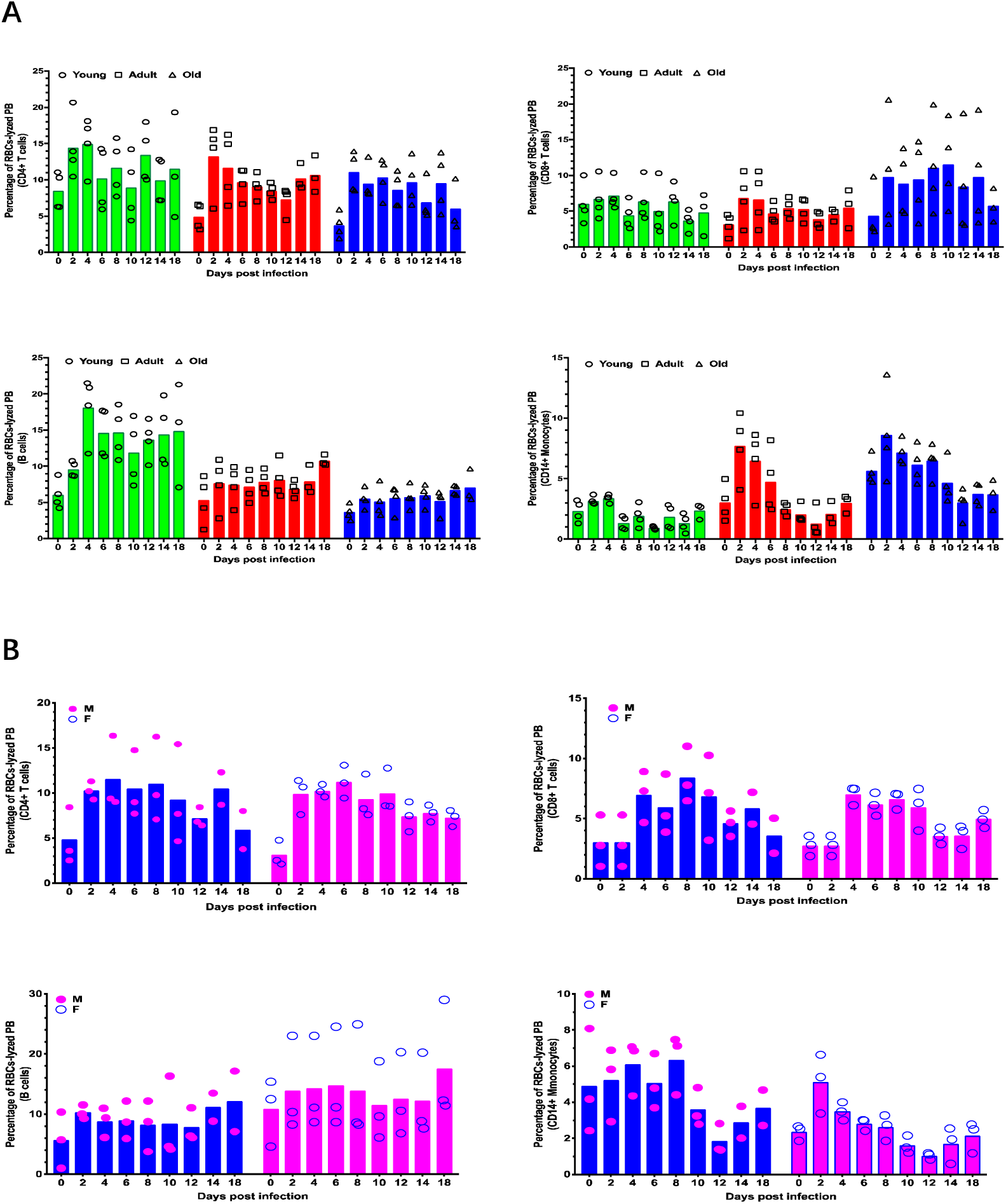
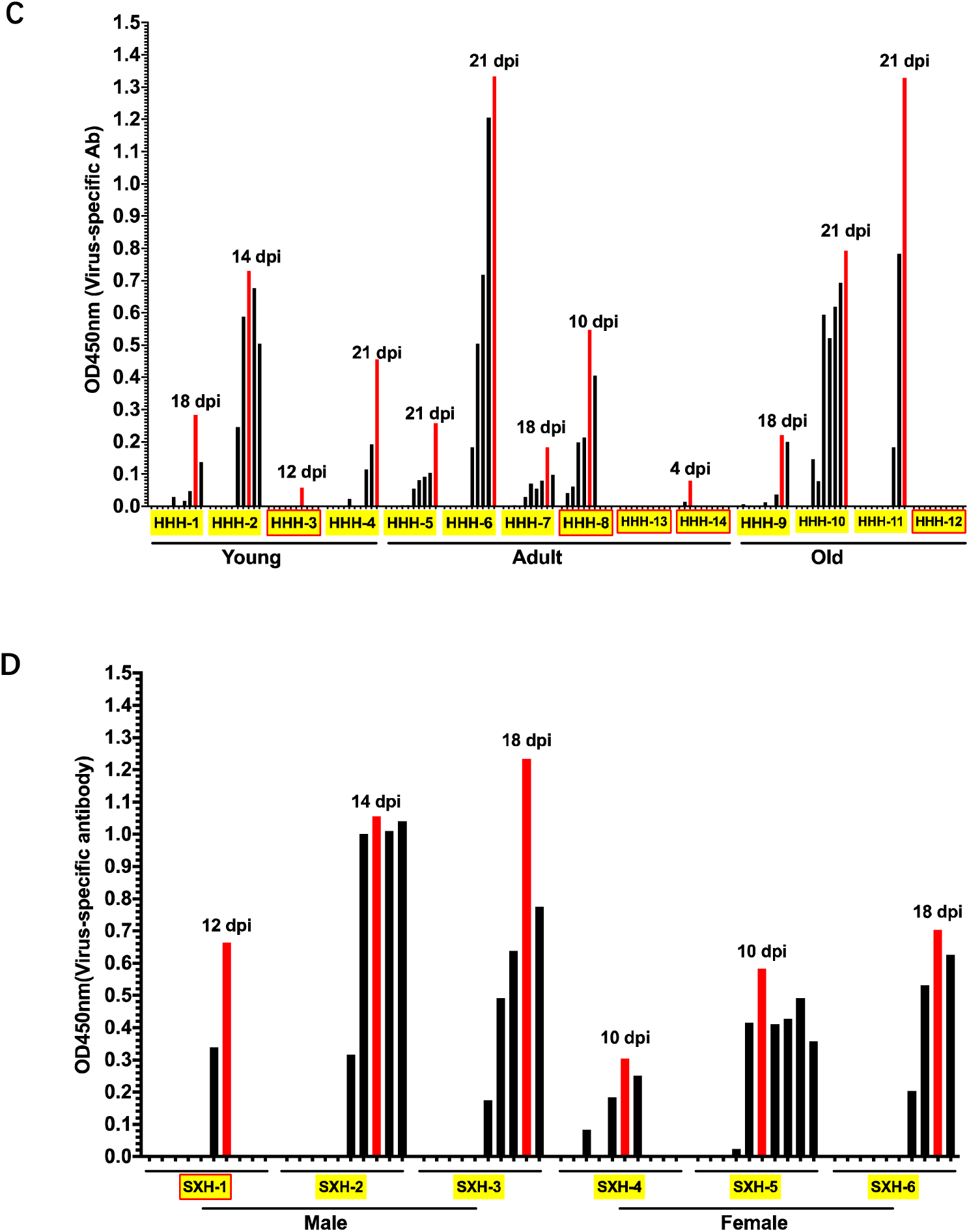
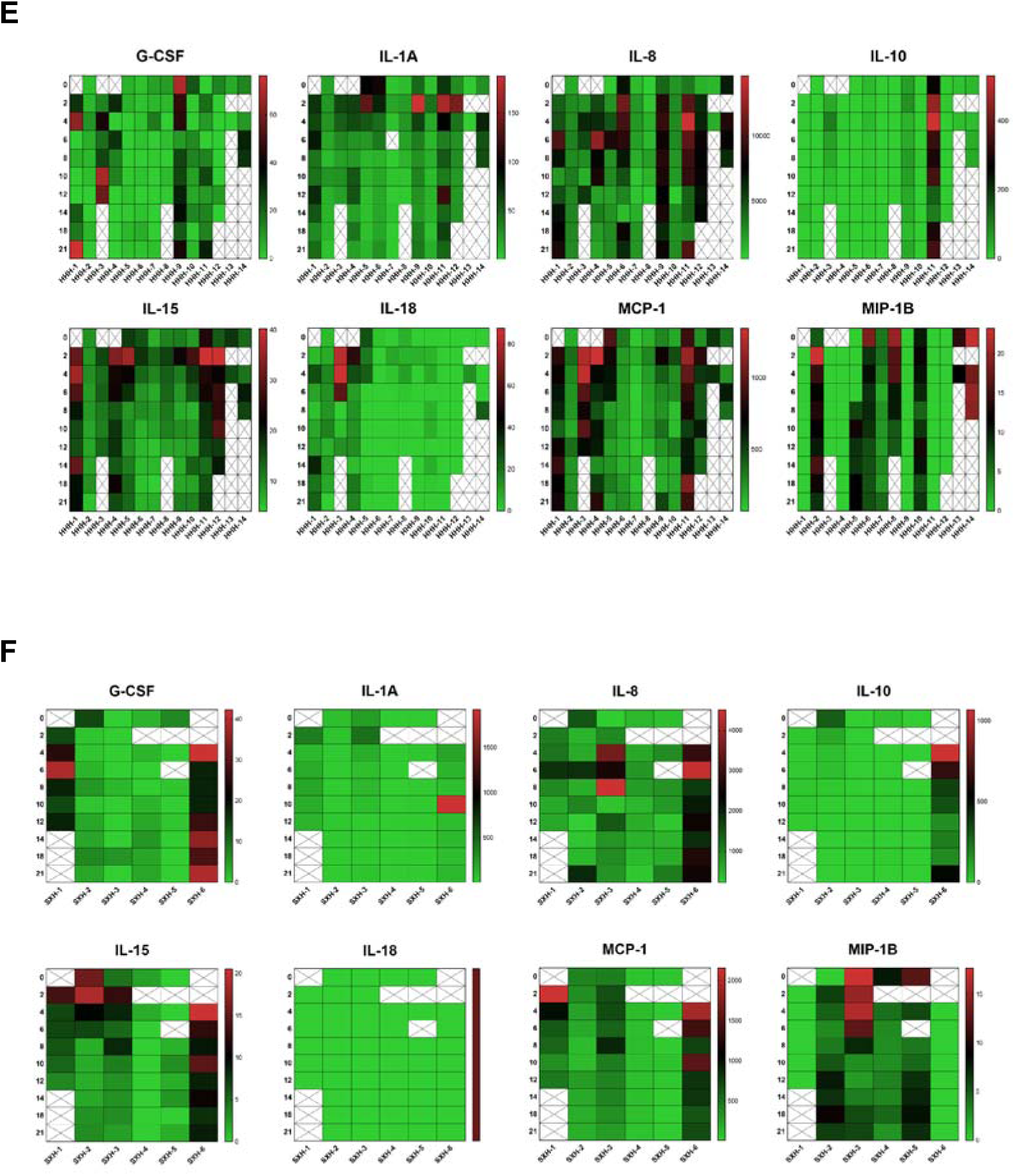
Host responses to SARS-CoV-2 infection. Responses of immune cells to viral infection were measured by flow cytometric analysis of cellular subpopulations in peripheral blood of infected *M. mulatta* (A) and *M. fascicularis* (B). Antibody responses of *M. mulatta* (C) and *M. fascicularis* (D) to viral infection was evaluated by total virus-specific antibody via ELISA. Inflammatory cytokines in serum samples from *M. mulatta* (E) and *M. fascicularis* (F) were measured by Luminex Technology Multiplex Assays as described in Materials and Methods.

Humoral immune responses to SARS-CoV-2 infection were evaluated in viral S protein-coated ELISA plates. Virus-specific antibodies became detectable as early as 4 dpi in both species of old world monkeys. In the following week, some animals showed a peak level of virus-specific antibodies. In the other animals, antibody levels gradually increased until the end of experiment (21 dpi) (Figure 4C). On 30 dpi, we still observed that antibody levels in 2 *M. mulatta* (one adult HHH-6 and one old HHH-11) and one *M. fascicularis* (adult female SXH-6) kept increasing (data not shown). One of old *M. mulatta* hadn’t produce detectable virus-specific antibody within 21 days of experiment. However, young *M. mulatta* showed overall lower levels of virus-specific antibodies compared with those in adult and old *M. mulatta*. Adult *M. fascicularis* gave the comparable or slightly higher levels of antibodies than young *M. mulatta.* No difference of antibody level was noted between SARS-CoV-2 infected male and female *M. fascicularis* (Figure 4D). We couldn’t detect virus-specific antibodies in serum samples from infected *C. jacchus* probably due to lack of cross-reactivity of antibody detection kit with samples from *C. jacchus.*

Furthermore, host inflammatory responses to SARS-CoV-2 infection were assessed via non-human primate multiplex inflammatory cytokines assay. Twenty-three inflammatory cytokines in serum samples from *M. mulatta* and *M. fascicularis* were quantitated and dynamically analyzed at the indicated time points. We found that eight cytokines (G-CSF, IL-1A, IL-8, IL-15, IL-18, MCP-1, MIP-1B, sCD40-L), among 23 cytokines assayed, were detected in most of animals. Especially, expression of sCD40-Lwas beyond the upper limitation of detection in most of tested samples (data not shown). IL-10 was only induced in monkeys of HHH-11 and SXH-6 on 4 dpi. In addition, IL-1A, IL-8, IL-15 and MCP-1 were strikingly induced in HHH-11 and SXH-6 and peaked on 2 or 4 dpi. Monkey HHH3 produced a peak of IL-15, IL-18 and MCP-1 on 2 dpi after SARS-CoV-2 infection, followed by gradual decline. Second peaks of some cytokines occurred at the late stage of experiment (Figure 4E & 4F).

Finally, in order to further investigate histopathogical responses to SARS-CoV-2 infection, we necropsied 8 animals on 4 dpi (HHH-13), 7 dpi (HHH-14), 12 dpi (HHH-8), 13 dpi (SXH-1, RH-2, RH-5), 15 dpi (HHH-3 and HHH-12). Severe gross lesions were observed on lung, heart and stomach of *M. mulatta* and *M. fascicularis*, but not *C. jacchus*. The main gross lesions involved massive pulmonary punctate haemorrhage, swollen Hilar and mediastinal lymph nodes, pericardial effusion, and swollen Mesenteric lymph nodes (Figure 5A). Histopathological analysis showed that thickening of the pulmonary septum and infiltration of inflammatory cells, diffuse hemorrhage and necrosis in *M. mulatta*. Exudate, a small number of lymphocytes and neutrophils were found in some blood vessels of *M. mulatta* liver, and some hepatocytes were swollen, with multiple vacuole like cells (mild) steatosis. The structure of spleen was clear, the number of germinal centers increased significantly, and capillary hemorrhage occurred in paracortical area. The number of germinal centers of mesenteric lymph nodes significantly increased. The space of renal vesicle space was widened and some glomeruli were atrophied. In *M. fascicularis*, histopathological changes in the respiratory system were similar to those examined in *M. mulatta*. Hyperplasia of mesenteric lymph nodes was observed in *M. fascicularis*. Mild infiltration of local inflammatory cells was also examined in the kidney of infected *M. fascicularis. C. jacchus* showed slight infiltration of inflammatory cells into the broken pulmonary septum with some necrotic cells. The hepatocytes were swollen. Light hemorrhage was observed in spleen, the germinal center of which is in actively proliferative condition (Figure 5B). Furthermore, ultrastructural examination under transmission electronic microscope was conducted on 70-80nm ultrathin sections. Although we didn’t find viral particles in any of examined tissues, typical ultrastructural lesions were observed in lung (increased type II pneumocytes, red blood cells and macrophages in alveolar airspace) and heart (increased number of mitochondria in the myocardium, loose arrangement of muscle fibers and fracture of some mitochondrial ridges) (Figure 5C), which was consistent to the results from microscopic inspection.

**Figure 5.**
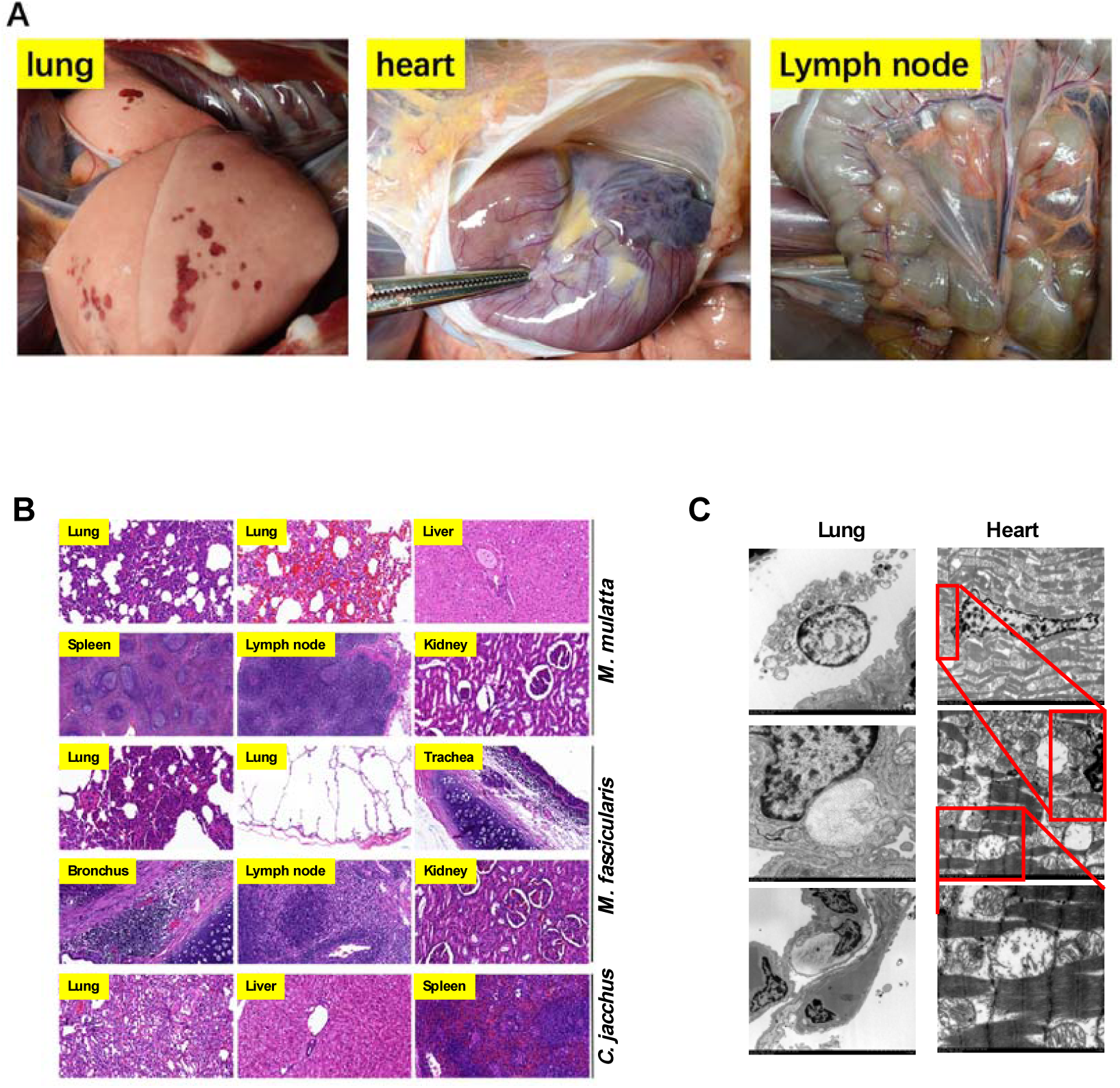
Pathological evaluation of tissues post SARS-CoV-2 infection. At necropsy, gross lesions were examined and recorded. Representative gross lesions of lung and lymph nodes were shown in panel (A). Tissue samples were cut into proper size and fixed in 10% neutral buffered formalin for HE staining, followed by microscopic inspection (B). Tissue samples of lung (left) and heart (right) fixed in 2.5% glutaraldehyde and 1% osmium tetroxide solution were for ultrastructural analysis (C). Histopathological changes were described in the text.

## DISCUSSIONS

SARS-CoV-2, recently reported as another novel coronavirus escaping from Pandora’s box after16 years ^13^, leaves us not just many mysteries but also hopes in its jar. Animal models of COVID-19, particularly non-human primate models, will give us hopes to uncover many mysteries of SARS-CoV-2 and COVID-19. Since December of 2019, scientists all over the world have been working to establish animal model of COVID-19. So far there are several models reported in preprinted journals or peer-reviewed published journals, including two small murine models (hACE2 transgenic mouse^14^ and Gold Syrian hamster^15^), one ferret model^16^ and four non-human primate models (three *M. mulattta* models^5-7^, one *M. fascicularis* model^17^). Due to differences of viral isolates, challenging routes, and animal species, controversial results or conclusions were drawn in these reported models of COVID-19. In this study, our comparative study of SARS-CoV-2 infection demonstrates that *M. mulatta* is the most susceptible animal to viral infection among 3 species of non-human primates, followed by *M. fascicularis* and *C. jacchus*. Age and gender may not affect the infection of SARS-CoV-2 in these monkeys. The non-human primate model in this study recapitulates several clinical features of COVID-19.

Retrospective studies of clinical features of COVID-19 show that older patients have more severe diseases, which may be associated with comorbidities, including tumor, diabetes, obesity, hypertension, heart diseases and so on^18-20^. These underlying diseases may make senior patients more susceptible to SARS-CoV-2 infection and worsen COVID-19. However, animals in this study were screened prior to experiments and supposed to be no severe complications/comorbidities and relatively healthy with normal immunity. This may explain that no matter age and gender within one species, monkeys showed similar susceptibility and responses to SARS-CoV-2 infection.

Monitoring body temperature is one of critical steps to define COVID-19 suspected patients, which leads to the conclusion that fever is the most common in COVID-19 patients, counting for more than 85% of all inpatients^3,18,21^. Here, all SARS-CoV-2 inoculated *M. mulatta* had increased body temperature at some time points with a peak of 40.9 °C. One-third of *M. fascicularis* and *C. jacchus* had a slightly elevated body temperature. Nevertheless, this manifestation could not be defined as fever since we don’t have the physiological body temperature of various species of monkeys as a reference.

The other important clinical feature of COVID-19 is abnormal changes of chest radiograph from X-ray and CT^22^, which is generally considered to be direct evidence for pneumonia. Ground glass opacities (GGO) were seen in 40%-70-89.6% of COVID-19 patients, followed by consolidation (13-33.9%) and other abnormal radiographs^18,21,23,24^. In this study, all animals (*C. jacchus* not determined) began to show progressively abnormal chest radiograph (Figure 2B and Supplemental Figure 1) from the 10^th^ day after virus inoculation, which simulates COVID-19 to some extent. In order to confirm results of chest radiographs, we played a very important match game and found that in some cases, gross lesions on lungs observed after necropsy strikingly matched abnormal spots of chest radiographs collected right before necropsy.

Laboratory routine assays include BCT, BCA, biochemistry indexes, flow cytometric analysis, results of which could be used to evaluate patients’ health and immune status. More than 50% COVID-19 patients have lymphopenia and/or esoinopenia, indicating their immunity may be affected by viral infection^3,18,25^. High levels of D-dimer, CRP, procalcitonin and ALT are assayed in severe patients, which suggests abnormal functions of liver and blood coagulation^26-28^. By flow cytometric analysis of peripheral blood from *M. mulatta* and *M. fascicularis* in the present study, we showed the changes and trends of T cells, B cells and monocytes after viral infection (Figure 4B), indicating that host immune responses were induced by SARS-CoV-2 infection^29^. However, our results from the other laboratory assays did not correlate with clinical indexes.

Detection of viral genomic RNA is the most important evidence for diagnosis of SARS-CoV-2 infection. Dynamic analysis of virus shedding will be beneficial for treatment and control of COVID-19. Comparative studies of large amount of clinical samples from SARS-CoV-2 infected patients demonstrate that 93% bronchoalveolar lavage fluid samples were viral RNA positive, followed by sputum (72%), nasal swabs (63%), feces (29%) and blood/serum (1%)^30^. In samples from upper respiratory track, nasal swabs had higher levels of viral RNA than those in throat swabs^30^. SARS-CoV-2 RNA in nasal swabs was detectable at the onset of disease, and peaked on day 2 post onset of disease. After that, it began to decline slowly, and usually last for about 2 weeks^31^. In the present study, dynamics of virus shedding in the NHP was similar to that reported in COVID-19 patients, although some minor disparities existed due to differences between disease onset in patients and viral inoculation time in animals. On day 2 post viral inoculation, there was an unusual peak of viral genome RNA in nasal swabs (Figure 3A). We speculated that this may be the left-over of viral nucleic acid from viral inoculation, but not from replicating viruses since it sharply declined on 4 dpi. The real peak of detectable RNA from replicating viruses should be from 6 dpi to 8 dpi in nasal swabs from infected animals. In addition, detectable viral RNA in anal swabs and expression of ACE2 in digestive tract of patients with covid-19 suggest that fecal-oral routine has a great potential risk in the transmission of SARS-CoV-2^32^. In this study, we noticed that high level of virus shedding via feces lasted for a long time in 4 monkeys of animal model (Figure 3A).

Viral load in various tissues, as an indicator of tropism, correlates with viral pathogenesis. Lung is widely considered to be the target organ for SARS-Cov-2 replication since plenty of viruses are detected in bronchoalveolar lavage fluid from COVID-19 patients^30^. At the earlier stage of our animal models, we could detect SARS-CoV-2 viral genomic RNA in lung and the other respiratory tissues (Figure 3C). However, SARS-CoV-2 RNA became negative in lung after 10 dpi, but was still detectable in weasand, and bronchus. It is possible that inflammatory environment of lung may push viruses out to other tissues.

Appearance of viral RNA in blood, out of lung, correlated with the severity COVID-19^33^. We could detect viral RNA in blood samples from all of 24 monkeys, although there were some differences of viral load and lasting time among individual animals. However, we couldn’t find the correlation between blood viral RNA and severity of disease.

Finally, histopathology is the golden standard and very important for evaluation of COVID-19 patients. So far, most of the clinical reports on the pathological changes of covid-19 come from the aged patients with comorbidities, focusing on lungs. Typical histopathology of pneumonia is observed in COVID-19 patients in additional to mild inflammation in liver and heart ^34^. After SARS-CoV-2 infection, *M. mulatta* and *M. fascicularis*, but not *C. jacchus*, showed severe histopathological changes in lung as pneumonia, and inflammation in liver and heart.

In conclusion, the NHP model in this study simulates several important aspects of COVID-19. Using this model, we should further explore pathogenesis of SARS-CoV-2 in order to answer some critical remaining questions in clinics, such as the refractory patients, cytokine storm, antibodies against SARS-CoV-2, and so on. And this model is suitable for preclinical evaluation of anti-viral drugs and vaccines against SARS-CoV-2.

## METERIALS AND METHODS

### Ethics statement

All animal procedures were approved by the Institutional Animal Care and Use Committee of Institute of Medical Biology, Chinese Academy of Medical Science (Ethics number: DWSP202002 001), and performed in the ABSL-4 facility of Kunming National High-level Biosafety Primate Research Center, Yunnan China.

### Virus amplification and identification

Viral stock of SARS-CoV-2 was obtained from the Center of Diseases Control, Guangdong Province China. Viruses were amplified on Vero-E6 cells and concentrated by ultrafilter system via 300kDa module (Millipore). Amplified SARS-CoV-2 were confirmed via RT-PCR, sequencing and transmission electronic microscopy, and titrated via plaque assay (10^6^ pfu/ml).

### Animals and experimental procedures

Three species of monkeys from two families of primates (old world monkeys and new world monkey) were used for this study. Detailed information about experimental animals was shown in Figure 1A.

Animal groups and experimental schedules were outlined in Figure 1A. Referring to the NHP model of SARS, we inoculated old world monkeys with total 4.75ml of 10^6^ pfu/ml SARS-CoV-2 intratracheally (4ml), intranasally (0.5ml) and on conjunctiva (0.25ml), new world monkeys with 1.0ml intranasally. Animals were daily checked for clinical sign and body temperature. At the indicated time points in Figure 1A, we anaesthetized animals with ketamine and performed the following experimental procedures. Every other day post inoculation, we took chest radiography of animals, harvested peripheral blood to prepare samples of whole blood or serum, and collected nasal, pharyngeal, and rectal swabs in 800ul Trizol LS solution (Invitrogen, US) for further analysis. Monkeys with severe signs were chosen for necropsy and pathological changes of all organs were recorded and evaluated at gross, histological and ultrastructural levels.

### Chest radiograph

Chest X-ray image of each anaesthetized monkey was taken with 55-75v and 8-12.5mA every other day using Mobile digital medical X-ray photography system (MobileCooper, Browiner China). Data was evaluated and scored double-blindly and independently by two radiologists.

### Quantification of viral genome in swabs, feces or tissue samples

400 ul of Trizol LS-treated and inactivated samples was used for RNA extraction according to the Direct-zol™ RNA MiniPrep protocol (ZYMO RESEARCH CORP, US). 50 μl of DNase/RNase-Free Water was used to elute RNA. Real time RT-PCR was used to quantify viral genome in samples using TaqMan Fast Virus 1-Step Master Mix (ThermoFisher, US) and purified viral RNA of SARS-CoV-2 as a standard curve, performed on CFX384 Touch Real-Time PCR Detection System (Biorad, US). Conditions for RT-PCR were used as follows: 25°C for 2min, 50°C for 15min, 95°C for 2min, then 40 cycles at 95°C 5sec and 60°C 31sec. Primers and probe, specific for *NP* gene was synthesized according to sequences reported by China CDC, Target-2-F:

GGGGAACTTCTCCTGCTAGAAT, Target-2-R:

CAGACATTTTGCTCTCAAGCTG, Target-2-P:5’-FAM-

TTGCTGCTGCTTGACAGATT-TAMRA-3’

### Hematology

Blood was collected in the tube containing anti-coagulant from inguinal veins of anaesthetized animals for Blood Clotting Test (BCT), Complete Blood Count (CBC) and flow cytometric analysis. BCT was conducted according to the manufacture-recommended protocol (Sysmex CA-600, Japan). Items in BCT includes Fbg sec, Fgb C, PT%, PI-INR, PT-R, PT-T, TT-T and APTT-T. CBC was performed on Auto Hematology Analyzer (BC-5000 Vet, Mindray, China), including white blood cells-related (numbers and percentages of Neu, Lym, Mon, Eos, Base), red blood cells-related (RGB, HCT, MCV, MCH, MCHC, RDW-CV) and platelet-related items (PLT, MPV, PDW, PCT). Flow cytometric analysis was conducted on Cytoflex (Backman, USA) using florescent-labelled antibodies against cellular surface marker CD45, CD3, CD4, CD8, CD14. Briefly, 50 ul of whole blood was incubated with proper fluorescent-labelled antibodies for 30min in dark at room temperature. 450 ul of 1x RBC Lys solution (BD) was added to the antibody-blood mixture and incubated for 15min at room temperature to remove red blood cells, which was ready for analysis on Cytoflex machine.

### Blood biochemical indexes

Non anti-coagulant blood from inguinal veins of anaesthetized animals was collected for serum preparation. Biochemical indexes in serum were measured on the automatic biochemistry analyzer (KHB ZY-1280, Shanghai Kehua Bio-engineering, China) using commercial kits, including liver function-related (TP, ALB, GLB, A/G, TB, DB, IBIL, ALT, AST, ALP), kidney function-related (BUN, UA, Cr), blood lipid-related (TC, TG, HDL, LDL, AP0A1, AP0B, LP(a)), heart muscle-related (CRE-E, LDH, α-HBDH, CK, CK-MB), humeral immunity-related (IgM, IgG, IgA, C3 and C4), and others indexes (GLU, CRP).

### Histological evaluation

For paraffin-bedded sections, tissues were collected and fixed in 10% neutral-buffered formalin, embedded in paraffin. 5 um sections were prepared for haematoxylin and eosin (H&E) staining, RNAscope or immunofluorescent staining. The whole H&E slide was scanned via 3D HISTECH system, evaluated by a pathologist double-blindly.

For ultrastructural sections, tissues were successively fixed in 2.5% glutaraldehyde and 1% osmium tetroxide solution. After gradual dehydration in various concentrations of ethanol, tissues were embedded in epoxy resin and cut into 60-70nm sections. After stained with 2% uranium acetate, ultrathin sections were examined under Hitachi transmission electron microscope H-7650.

### Localization of viral RNA in tissues

A novel nucleic acid hybridization technique RNAscope was performed to detect and localize viral RNA in paraffin-embedded tissue sections using SARS-CoV-2 specific probe (RNAscope® Probe-V-nCoV2019-S, ACD, Cat No. 848561, targeting the region of 21631 – 23303 (NC_045512.2) of SARS-CoV-2 genome). FFPE slides were pretreated by baking, de-paraffinizing, target retrieval and protease treatment. SARS-CoV-2 specific probe was applied to the pretreated slides and hybridized with viral genome. Fluorophore was added to detect the hybridization on slides, followed by counter-staining with DAPI. Slides were scanned via 3D HISTECH system for further analysis.

### Virus-specific antibody response

Levels of SARS-CoV-2-specific antibodies in serum samples were evaluated via the commercially available SARS-CoV-2 Antibody Assay Kit (ELISA) (Cat#XG100H2, China). In this kit, viral spike protein was coated in each well of 96-well plate to capture spike-specific antibodies. HRP-conjugated goat anti-human IgG (H+L) antibody was added to detect the captured spike-specific antibodies. Data was plotted via the software GraphPad.

### Multiplex analysis of cytokines in serum

MILLIPLEX MAP Non-Human Primate Cytokine Magnetic Bead Panel - Immunology Multiplex Assay (PRCYTOMAG-40K, Millipore USA) were used in this study to assay 23 inflammatory cytokines according to the manufacturer’s protocol, which was performed on Bio-plex machine. Inflammatory cytokines in this panel included IL-1β, IL-4, IL-5, IL-6,IL-8/CXCL8,G-CSF,GM-CSF,IFN-γ,IL-1ra,IL-2,IL-10,IL-12p40,IL-13,IL-15,IL-17A/CTLA8,MCP-1/CCL2,MIP-1β/CCL4,MIP-1α/CCL3,sCD40L,TGF-α,TNF-α,VEGF-A and IL-18.

## ACKNOWLEDGEMENTS

The authors would like to thank all staffs in National Kunming High-level Biosafety Primate Research Center for providing ABSL3- and ABSL4-related services. This study was supported by 2020YFC0841100 and 2020YFC0846400.

## AUTHOR CONTRIBUTIONS

SY, HL and XP designed the study; SY, YZ, WH, YY, JG, and HL wrote the manuscript; SY, YZ, WH, YY, JG, JW, DK, HW did the animal experiments.; MY, CM, SZ, JL, YD detected viral RNA; JX did RNAscope; LJ and XL performed immunofluorescent assay; HL, SY, YY, JY and YZ analyzed data; WY and HL detected cytokines and antibodies. XQ YL, and FL prepared Vero-E6 cells; KD, DW, HL and YZ did histopathological work; HZ and CK provided viral stock; QS, YH, QD, QL gave suggestions to the study. All authors have approved the submitted manuscript.

